# Study and Modeling of Biological Noise-Filtering Properties of Conserved Gene Regulatory Networks Motifs in Animal Development

**DOI:** 10.1101/2022.06.13.495911

**Authors:** Lina M. Ruiz G., Machado R Gloria, Boris A. Rodriguez R

## Abstract

Biological noise results from heterogeneous gene expression levels among a group of cells [1]. This heterogeneity is due to the variation in gene expression that occurs over time at the single-cell level. Some noise-filtering mechanisms like redundancy in genetic circuits have been identified. Likewise, the feed-forward loop network motif has been found to have noise-filtering capacities in animal development. On the other hand, previous studies have contradictory conclusions about the noise-filtering capacities of the feedback loop and none of them have studied this capacity in the activator-inhibitor regulatory system. Here we studied some dynamical properties, such as noise and expression levels, in self-activated and activator-inhibitor regulatory systems, both at the unicellular and multicellular levels. These systems are essential in the self-patterning and community effect processes occurring in development and differentiation. We used the three-stage model to represent the expression of a gene with promoter regulation and Hill functions to represent the regulatory connections between genes. We used Gillespie’s Algorithm and the Chemical Langevin Equation for simulations. The regulatory systems evaluated do not reduce the biological noise. On the contrary, the noise remains at the same level or increases in comparison with an unregulated gene. The noise levels in these systems depend on the gene expression type of both the regulator and the regulated gene. In this way, the particular forms in which genes connect to each other in these regulatory systems do not explain the noise in expression. However, the noise has a propagation pattern different for activation and inactivation types of regulation. Finally, the diffusion and colony size could be mechanisms of noise filtering in gene expression in a colony of cells. The increase in diffusion rate and colony size are necessary to synchronize gene expression and perform the community effect in embryonic development.

## 1. INTRODUCTION

Biological noise results from heterogeneous gene expression levels among a group of cells [1]. This heterogeneity is due to the variation in gene expression that occurs over time at the single-cell level [2]. There are two different sources of biological noise, *extrinsic* and *intrinsic* [3]. Intrinsic sources have greater influence than extrinsic sources in determining noise, and they play a significant role when expression regulation happens at the promoter level [4–6]. Regulation at the promoter level generates transitions between an *active state* (ON), in which mRNA is transcribed, and an *inactive state* (OFF), in which mRNA is not transcribed [7–9]. As a consequence, transcription commonly occurs discontinuously over time [10,11]. This means that most genes are transcribed in short periods called *transcriptional bursts*, interspersed by silent intervals [2,12,13]. T*ranscriptional bursting* has been observed across many species and it is one of the main sources of noise [10,11].

Transcriptional burst expression has been reported in some genes involved in development, like the *hindsight* (hnt) and *u-shaped* (ush) genes in *Drosophila* embryos [14]. It has also been observed in the gap gene *hunchback* in *Drosophila* when it is expressed in the anterior pole of the embryo [15]. In mouse embryonic stem cells, the *nanog* gene shows transcriptional burst kinetics [15]. Most of the developmental regulators are thought to be transcribed in bursts [15].

Sometimes, noise is useful because it drives phenotypic variability in developmental transitions [10,15,16]. For example, this is the case with neural fate, which is stochastically determined [17]. But other times, embryos must establish and maintain precise levels of gene expression [18]. This is the case with the process of differentiation and patterning formation, in which many mechanisms that buffer against expression noise have been observed [19,20]. One of such mechanisms is redundancy in genetic circuits. [21]. Shadow enhancers, which filter the noise in Transcription Factors (TF) by separating inputs (i.e., each enhancer responds to a different TF), are another example [21]. Likewise, Bone Morphogenetic Protein modulates the kinetic of the transcriptional burst of its target genes, reducing noise for some of them [14].

Noise-filtering has also been studied in gene regulatory network (GRN) systems found in the self-patterning and differentiation process in development. For instance, interlinked feed-forward loops have been found to be effective in filtering noise [16]. Two other systems, the self-activated and activator-inhibitor systems are also present in this developmental process. There are several studies that point to different conclusions regarding the noise-filtering capacities of the self-activated regulatory system. For instance, in [16], the feedback loop is found to be a noise controller. On the contrary, [22] report that noise is amplified in a positive feedback loop. To our knowledge, there are no studies regarding noise-filtering capacities for the activator-inhibitor system.

The self-activated and activator inhibitor systems play an important role in development. For instance, self-activated genes specify spatial domains of gene expression. Usually, new regulatory states are locked down by the deployment of a paracrine signal with positive feedback [23]. This means that the gene that encodes the paracrine signal responds to its own signal transduction system. Thus, all cells in the domain receive and emit the same signal. This activates a unique set of regulatory genes in a given spatial domain of an embryo [23]. This results in a phenomenon known as the “community effect” [23,24], in which the cells are linked together by intercellular signaling and by the expression of the same set of downstream genes [23,24].

The activator-inhibitor regulatory system is also very important for self-organized fate patterning [25]. It is a reaction-diffusion system in which two interacting and diffusing species of molecules can generate complex patterns [26]. It is made up of a short-range activator which enhances its production and the production of a long-range inhibitor [27]. One example of this are the ligands Nodal and Lefty. They constitute an activator-inhibitor system in animals as different as sea urchins and mice [28]. Nodal is required for the initial specification of the anteroposterior axis in mice, mesoderm formation in mice and *Xenopus*, left-right patterning in mice, and development of head and trunk mesoderm and endoderm tissues in zebrafish [28].

Here we studied some dynamical properties, such as noise and expression levels, in self-activated and activator-inhibitor regulatory systems, both at the unicellular and multicellular levels. This is a theoretical study in which dynamical properties are measured for changes in some kinetic parameters that affect the temporal dynamics of expression [9–11]. We varied those parameters within ranges that included values reported by experimental studies to answer the following questions: how do the dynamical properties of these regulatory systems vary in a cell throughout the range of parameters values? How does noise propagate from a regulator gene to a regulated gene? Could we describe these regulatory systems as noise-filtering mechanisms? What are the dynamical properties of a self-activated gene in a colony of cells?

## 2. METHODOLOGY

### 2.0 Generalities of models and simulations

In the regulatory systems evaluated here, the expression of each gene was represented with the three-stage model [29]. This model includes the activation and deactivation of the promoter, the synthesis and degradation of mRNA, and the synthesis and degradation of protein. This set of biochemical reactions was simulated with both Gillespie’s Algorithm (GA) and the Chemical Langevin Equation (CLE) for a unicellular and multicellular approach, respectively [30,31].

The GA is a scheme that simulates every reaction event and generates trajectories within the state probability distribution [32]. In each iteration of the algorithm, and according to the Propensity Functions (PF), both the type of reaction (i.e., activation, deactivation, synthesis, degradation, etc.) and the reaction time t are estimated. Each iteration of this algorithm produces very small changes in the system, so simulating the process for large systems is time-consuming [33]. This is why this algorithm was used for simulations at the unicellular level. Instead, to simulate the self-activated gene in a cell colony we used the CLE, since it represents the averaged dynamics after a set of these biochemical reactions occur in a time step [31]. To simulate the regulatory connections of the regulatory systems we used with Hill functions for activation and inactivation [34].

Noise is commonly measured with the Fano Factor (FF) and the squared coefficient of variation (CV^2^) of the gene’s temporal expression. The FF is the ratio between the variance and the mean [11]. The FF is key to quantifying how much a gene’s expression deviates from Poisson statistics, which is the characteristic behavior of a constitutive gene [35]. For a Poisson behavior, the FF equals one and it defines a “standard dispersion”. Therefore, distributions with an FF smaller or larger than one are considered under or over-dispersed, respectively [35].

Conversely, the CV^2^ measures the variability due to system size or molecule number [6]. CV^2^ is the ratio between the variance and the squared mean [6]. Noise measured with CV^2^ tends to increase when the size of the system decreases [36]. This is because changes in the number of molecules are more significant when the number of molecules is small than when it is larger. This phenomenon is known as the finite-number effect [36].

Changes in some kinetic parameters have been shown to affect noise and expression levels. These parameters are the promoter activation rate [37,38], the promoter deactivation rate [10], the mRNA degradation rate [9,10], and the protein degradation rate [10,11]. Here we estimated the dynamical properties (i.e., noise and expression levels) of self-activated and activator-inhibitor regulatory systems for a range of values of these parameters. Additionally, for a cell colony, we also evaluated the diffusion rate and the colony size, since these two parameters affect the amount of variability in gene expression levels between cells [23]. The main code of this study is in https://github.com/LinaMRuizG/BiologicalNoise_DevelopmentalGeneRegulatoryNetworks.

### 2.1 Noise and expression levels in an unregulated gene

We used the GA to simulate a gene without explicit regulation (from now on, referred to as an unregulated gene). For these genes, unlike for genes with explicit regulation (regulated genes from now on) the PF for promoter activation depends only on the activation rate (k_on_) and the promoter state (i.e. active -ON is 1-or inactive -OFF is 0-). The PF for the remaining biochemical reactions is the same for both regulated and unregulated genes. For instance, the promoter deactivation depends on the deactivation rate (k_off_) and the promoter state. The synthesis and degradation of mRNA depend on synthesis and degradation rates (ks_mRNA_ and kd_mRNA_), and the number of mRNA molecules (m). For the synthesis of mRNA we included a term that represents basal expression of mRNA by adding 0.01 to the PF. The synthesis and degradation of protein depends on synthesis and degradation rates (ks_p_ and kd_p_), and the number of protein molecules (p).

For unregulated genes, we simulated the system up to a final time of 1500 minutes (25 hours). We calculated the FF, CV^2^, and mean of molecules only for the steady-state. The first thirty percent of all data points for each time series (simulation) weren’t considered so that we could ensure that these estimators of dynamical properties were measured for the steady-state. We averaged these estimates over ten replicates and we plotted their values for each set of parameters. The simulations were made by changing the values of k_on_ and k_off_ between 0.01 and 1, and between 0.02 and 2, respectively. These parameters were selected based on previous studies [5,10,17]. The range for these parameter’s values was selected to include all the values found in experimental literature and previous models (Table S2-S8). However, there is a lack of knowledge about these kinetic parameters and many studies assume their own values [39–44]. The default values for all parameters are listed in Table S9.

We grouped the simulation results based on their similarity in dynamic behavior throughout the range of k_on_ and k_off_ values, using the estimates for FF, CV^2^, and mean of molecules. To do this, we randomly selected 500 pairs of values of k_on_ and k_off_ within their defined range. We estimated the average value of FF, CV^2^, and mean of molecules for 100 replicates of each pair. We then used the Gaussian Mixture Models clustering algorithm to find groups among the 500 pairs. This clustering was performed both at the mRNA and protein levels. The final group number we reported includes those found in both levels.

### 2.2 Noise and expression levels of the regulatory systems for individual cells

We used the GA to simulate the self-activated and activator-inhibitor regulatory systems in a single cell, which is equivalent to a set of decoupled cells. We compared the results with those of an unregulated gene (control). The PF for promoter activation depends on both the promoter state and k_on_. Unlike the case with unregulated genes, promoter activation for regulated genes also depends on a Hill function for each regulation type. For instance, the PF for self-activation, activation and inhibition in the activator-inhibitor system is calculated using the expressions 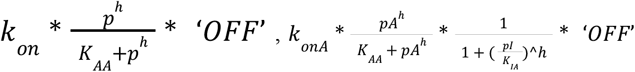 and 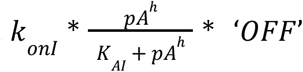, respectively, where p is the regulator protein, and K_**_ is the Michaelis-Menten constant, which represents the affinity of the *regulator protein* to the promoter sequence [34]. The PF for the other reactions is the same as those for an unregulated gene.

For regulated genes, we also simulated the system up to a final time of 1500 minutes (25 hours). We calculated the FF, CV^2^, and mean of molecules in the steady-state like we did for unregulated genes. We averaged these estimates over ten replicates and we plotted their values for each set of parameters. The simulations were made by changing the values of k_on_ and k_off_ between 0.01 to 1 and 0.02 to 2, respectively. The default values for all parameters are listed in Table S9.

We tested the significance of differences in FF, CV^2^, and mean of molecules between the two regulated gene systems (self-activated and activator-inhibitor) and the unregulated gene system. To do this, we randomly selected 500 pairs of k_on_ and k_off_ values within their defined range for each of the three systems. We estimated the average value of FF, CV^2^, and mean of molecules for 100 replicates of each pair for each system. We then performed a Mann-Whitney test for the sample of 100 replicates of each of the 500 pairs to compare each regulated system to the unregulated gene system.

### 2.3 Noise propagation from the regulator to the regulated gene

We used the GA to simulate activation and deactivation systems of regulation in one cell. We compared the results with those of an unregulated gene. For the activated gene, the PF for promoter activation is 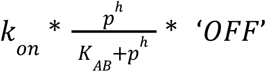, where *p* is the protein that activates the promoter (activator). For an inhibited gene, the PF for promoter activation is 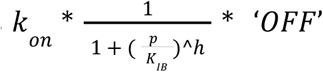, where *p* is the protein that inhibits the promoter (inhibitor). The PF for promoter activation of both the activator and inhibitor is k_on_*OFF, which is the same as that for an unregulated gene.

The PF for the other reactions are equal to those for an unregulated gene. We simulated and estimated FF, CV^2^, and mean of molecules throughout the range of k_on_ and k_off_ parameter values as previously described.

We performed two types of *in-silico* experiments. In the first one, the k_on_ and k_off_ parameters of regulators (i.e., activator and inhibitor) were changed, while those for regulated genes were kept unchanged. In the second one, the k_on_ and k_off_ parameters of regulated genes were changed, while those for regulators remained the same. We tested the significance of differences in FF, CV^2^, and mean of molecules between both systems and an unregulated gene system, as described in the previous section.

### 2.4 Noise and expression levels of the regulatory systems for coupled cells

We used the CLE to simulate a self-activated gene system in a cell colony. The colony is a population of isogenic cells arranged in a circular way that is similar to micropattern colonies [45]. From [31], we used the CLE implementation for burst production at the mRNA level but not at the protein level, since the default parameter satisfied that k_off_ >> kd_mRNA_ ∼ kd_p_ (Table S9). Additionally, we adapted this implementation for simulation in a cell colony. For this, each cell in the colony is expressing a self-activated gene which is represented by the set of CLE for mNRA (1) and protein (2):

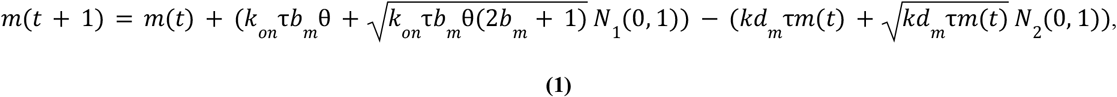

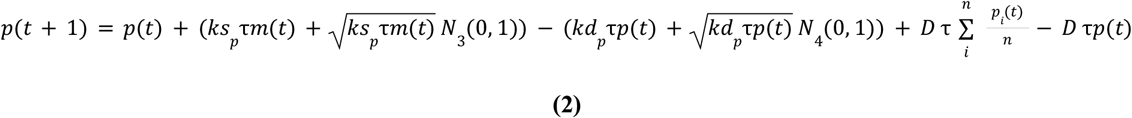

Where, k_on_ is the promoter activation rate, *b*_*m*_ is the mRNA burst size, θis the Hill function 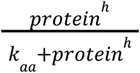 with *k*_*aa*_ equal to 100, N_i_ (0,1) is the white noise, kd_m_ is the mRNA degradation rate, ks_p_ is the translation rate, kd_p_ is the protein degradation rate, τ is the step size estimated with the *tau leap* (τ-leap) method as in [21], D is the diffusion rate, and n is the number of neighbors. The terms of diffusion were included in the CLE to couple the cells by their genetic product (i.e., a paracrine signal). These terms represent the diffusion into and out of each cell and its eight neighbors (i.e., Moore neighborhood). To avoid negative production values, we set a basal production value of 0.01 every time the result was below zero.

We simulated the system up to a final time of 1500 minutes (25 hours). We calculated the FF and mean of molecules for the whole colony in each time unit in the steady-state. We used the steady-state detection algorithm described in [46] to estimate the start of the steady-state. Then, we averaged the temporal FF and mean of molecules. The parameters we evaluated and their range of values are listed in Table 1. The default values for all parameters are listed in Table S9.

**Table 1.**
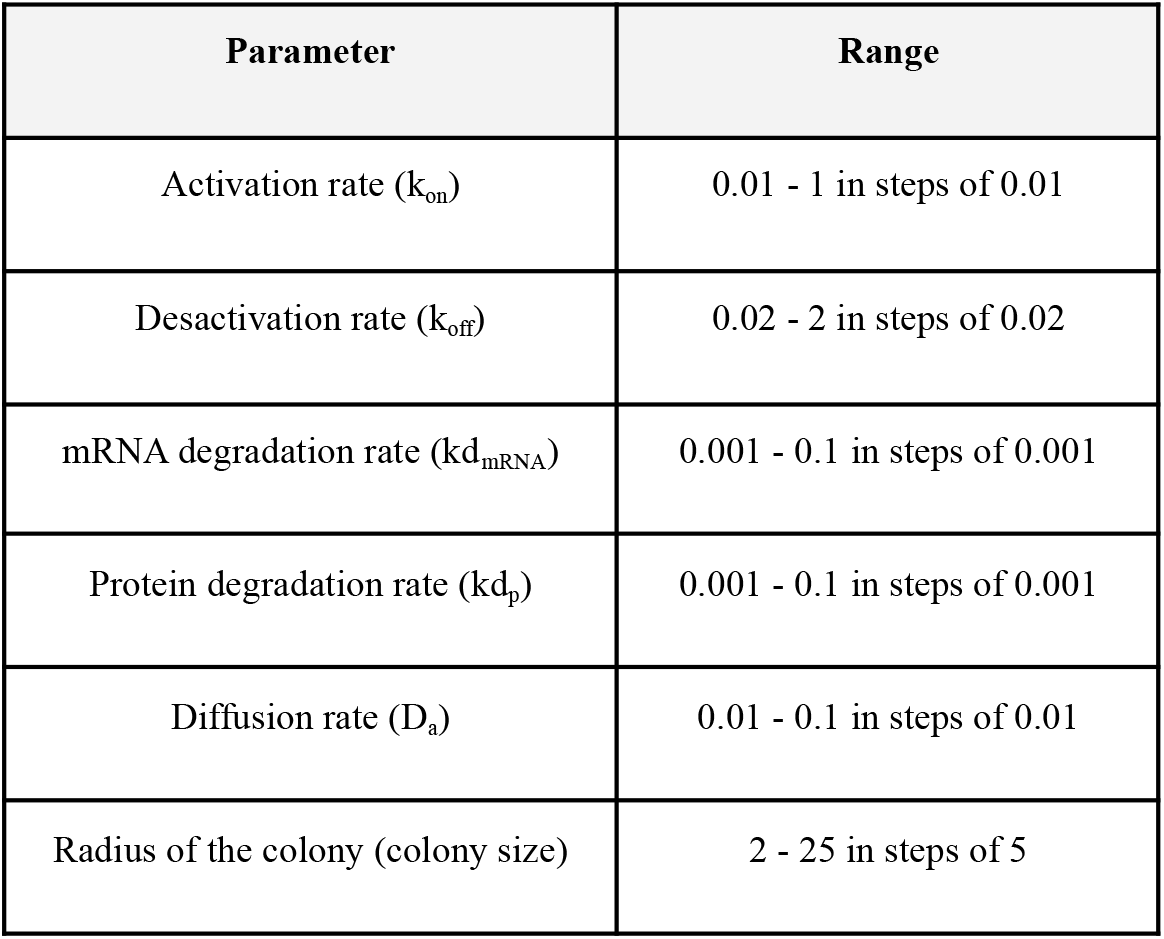
Parameter value ranges.

| Parameter | Range |
| --- | --- |
| Activation rate ($k_{on}$) | 0.01 - 1 in steps of 0.01 |
| Desactivation rate ($k_{off}$) | 0.02 - 2 in steps of 0.02 |
| mRNA degradation rate ($kd_{mRNA}$) | 0.001 - 0.1 in steps of 0.001 |
| Protein degradation rate ($kd_p$) | 0.001 - 0.1 in steps of 0.001 |
| Diffusion rate ($D_a$) | 0.01 - 0.1 in steps of 0.01 |
| Radius of the colony (colony size) | 2 - 25 in steps of 5 |

## 3. RESULTS

### 3.1 Noise and expression levels in an an unregulated gene

The estimated values for FF, CV^2^, and mean of molecules were qualitatively similar at both the mRNA and protein levels (Fig. 1 and S1). In both levels, there was an increase in the mean of molecules with the increase in k_on_, and there was a decrease in the mean of molecules with the increase in k_off_ (Fig. 1A and S1A). The noise measured with CV^2^ had an inverse relationship with the mean of molecules; larger systems had lower CV^2^ values (Fig. 1B and S1B). But there was no clear relationship between FF and mean of molecules, and the noise measured with FF had a different behavior from the noise measured with CV^2^ (Fig. 1C and S1C). Also at both the mRNA and protein levels, the FF had the highest values when k_on_ and k_off_ were in the area of low values, or in the Slow Promoter Kinetic (SPK) area [17,47]. In this area k_off_ could be larger than k_on_, k_on_ could be larger than k_off_, and k_off_ could be equal to k_on_.

**Figure 1.**
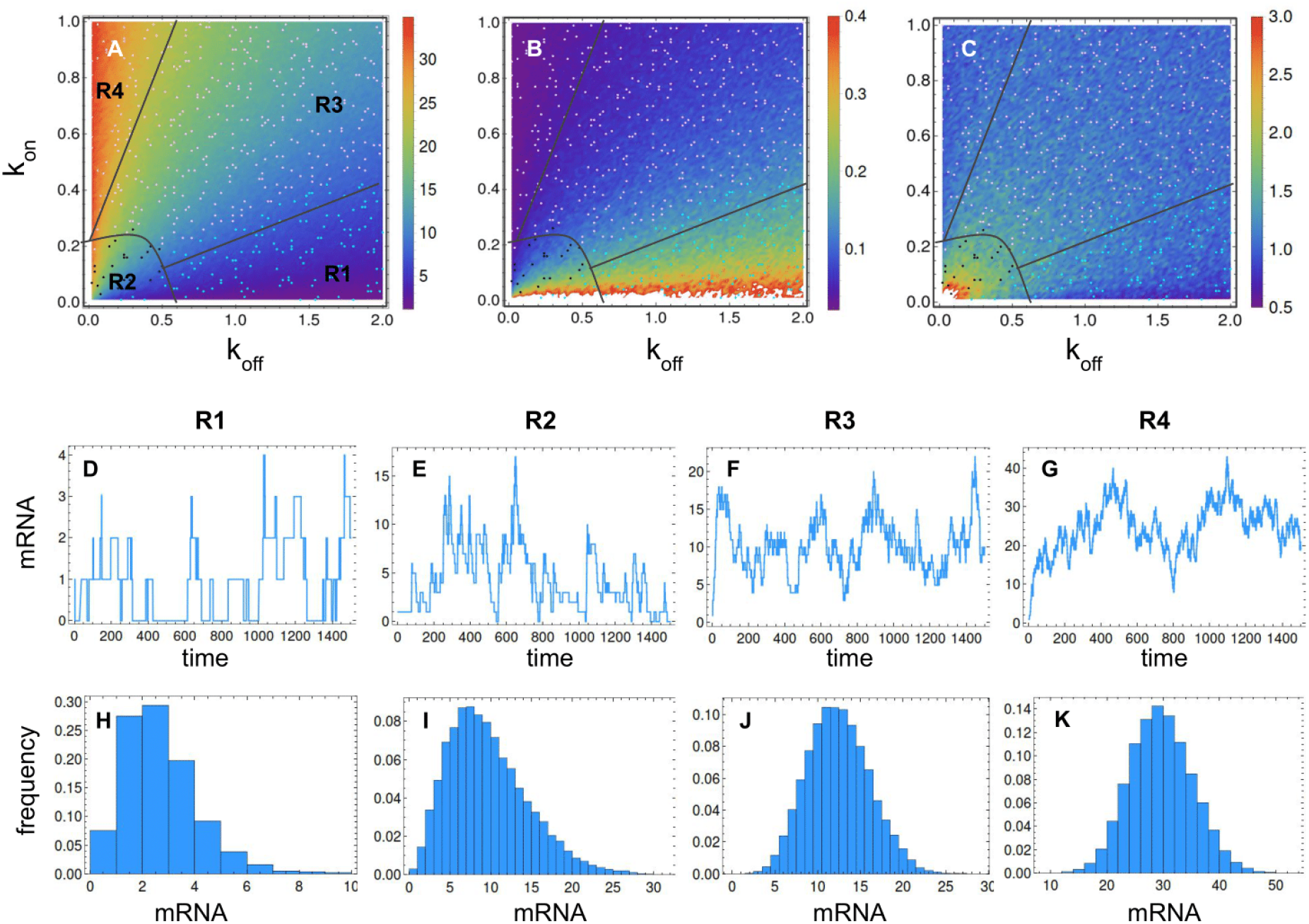
Regions of gene expression types at the mRNA level for an unregulated gene in a range of k_on_ and k_off_ parameter values. A. The four regions over the mean of molecules pattern B. The four regions over the CV^2^ pattern, C. The four regions over the FF pattern, D-G. The temporal dynamics of expression in each region, H-K. The steady-sate distribution in each region. Default parameters are listed in Table S9. The color of points represents the clustering group to which the pair of parameters belongs to. Color should be used for this figure in print.

The FF, CV^2^, and mean number of molecules that were estimated through the parameters k_on_ and k_off_ allowed us to identify four groups, or regions, of types of gene expression (Fig.1). This means that the expression dynamics of a gene regulated at the promoter level were similar in each region. In region 1, the promoter was mainly inactive because k_off_ was larger than k_on_ (Fig. 1A-C). As a result, the system was small and had the highest CV^2^ (Fig. 1A-B). Gene expression happened in discontinuous and small bursts with low variation, so FF had low values (Fig. 1D,H,C). In region 2, the promoter was in the SKP area, with low values for both k_off_ and k_on_ (Fig. 1A-C). Depending on the k_off_ and k_on_ ratio, the system was either small or large and had either a high or low CV^2^ (Fig. 1A-B), respectively. The expression happened in discontinuous bursts with high variation, as shown in the steady-state distribution, so FF had the highest values (Fig. 1E,I,C).

In region 3, the promoter had values outside of the SKP area for both k_off_ and k_on_ (Fig. 1A-C). Depending on the k_off_ and k_on_ ratio, the system was small or large and had high or low CV^2^ values (Fig. 1A-B), respectively. Gene expression happened in continuous bursts with low variation, as shown in the steady-state distribution, so FF was low (Fig. 1F,J,C). Finally in region 4, the promoter was mainly active because k_on_ was larger than k_off_ (Fig. 1A-C). Therefore, the system was large and had the lowest CV^2^ values (Fig. 1A-B). Expression was continuous with low variation, so FF was small (Fig. 1G,K,C). To summarize, throughout the regions gene expression went from happening in discontinuous bursts to being more continuous both at the mRNA and protein levels (Fig. 1-1S).

### 3.2 Noise and expression level of the regulatory systems for individual cells

There were non-significant differences in mean of molecules, CV^2^, and FF between an unregulated gene and a self-activated gene (A), both at the mRNA and protein levels (Fig. 2-S2 A-F). This means that the temporal dynamics of both genes were similar in all regions of expression (Fig. 1-S1). On the other hand, there were significant differences in mean of molecules, CV^2^, and FF between an unregulated gene and both genes A and I in an activator-inhibitor regulatory system for most regions except region 1, both at the mRNA and protein levels (Fig. 2-S2 A-C, G-L). In region 1, there were non-significant differences in FF between an unregulated gene and both genes A and I in an activator-inhibitor regulatory system, both at mRNA and protein levels (Fig. 2-S2 C, I, L).

**Figure 2.**
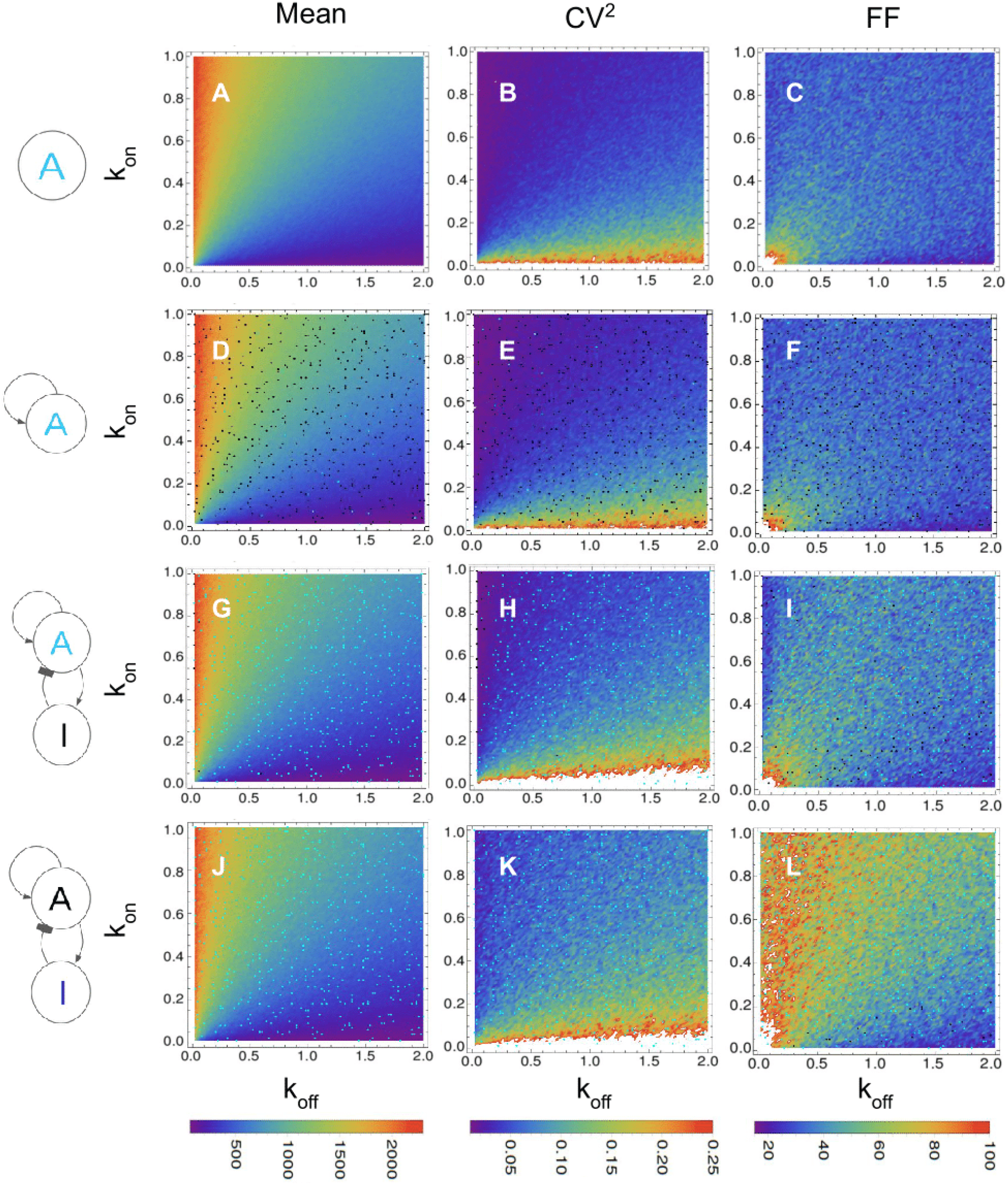
Estimates of mean of molecules, CV^2^, and FF at the protein level for gene expression of the regulatory systems evaluated in a range of values for k_on_ and k_off_. A-C. mean, CV^2^, and FF for an unregulated gene A, D-F. mean, CV^2^, and FF for a self-activated gene A, G-I. mean, CV^2^, and FF for gene A in an activator-inhibitor regulatory system when only the parameters of gene A were changed, J-L. mean, CV^2^, and FF for gene I in an activator-inhibitor regulatory system when only the parameters of gene I were changed. The black/cyan points indicate non-significant/significant differences when compared to an unregulated gene. The default parameters are found in Table S9. White points are values higher than the scale plotted. Color should be used for this figure in print.

### 3.3 Noise propagation from regulator to regulated gene

There were significant differences in FF, mean of molecules, and CV^2^ between an unregulated gene A and a regulated gene, both inhibited and activated, at the mRNA and protein levels (Fig. 3-S3). This was true for all regions, except for region 1, where there were non-significant differences in FF between an unregulated gene A and a regulated gene, both inhibited and activated, at the mRNA and protein levels (Fig. 3-S3 A and C). These differences imply higher values of FF and CV^2^, and lower values of mean of molecules for regulated genes compared to an unregulated gene. On the other hand, there was a similarity between the qualitative pattern of CV^2^ and the mean of molecules of regulated and unregulated genes (Fig. 2A-B, Fig.3 B-C and E-F), but the FF pattern was different (Fig. 2C, Fig.3 A and D). These differences in FF pattern depended on the type of expression. For instance, the inhibited gene had significant differences mainly in region 3, but it had non-significant differences in regions 1-2 and 4. In turn, the activated gene had significant differences in all regions except in region 1.

**Figure 3.**
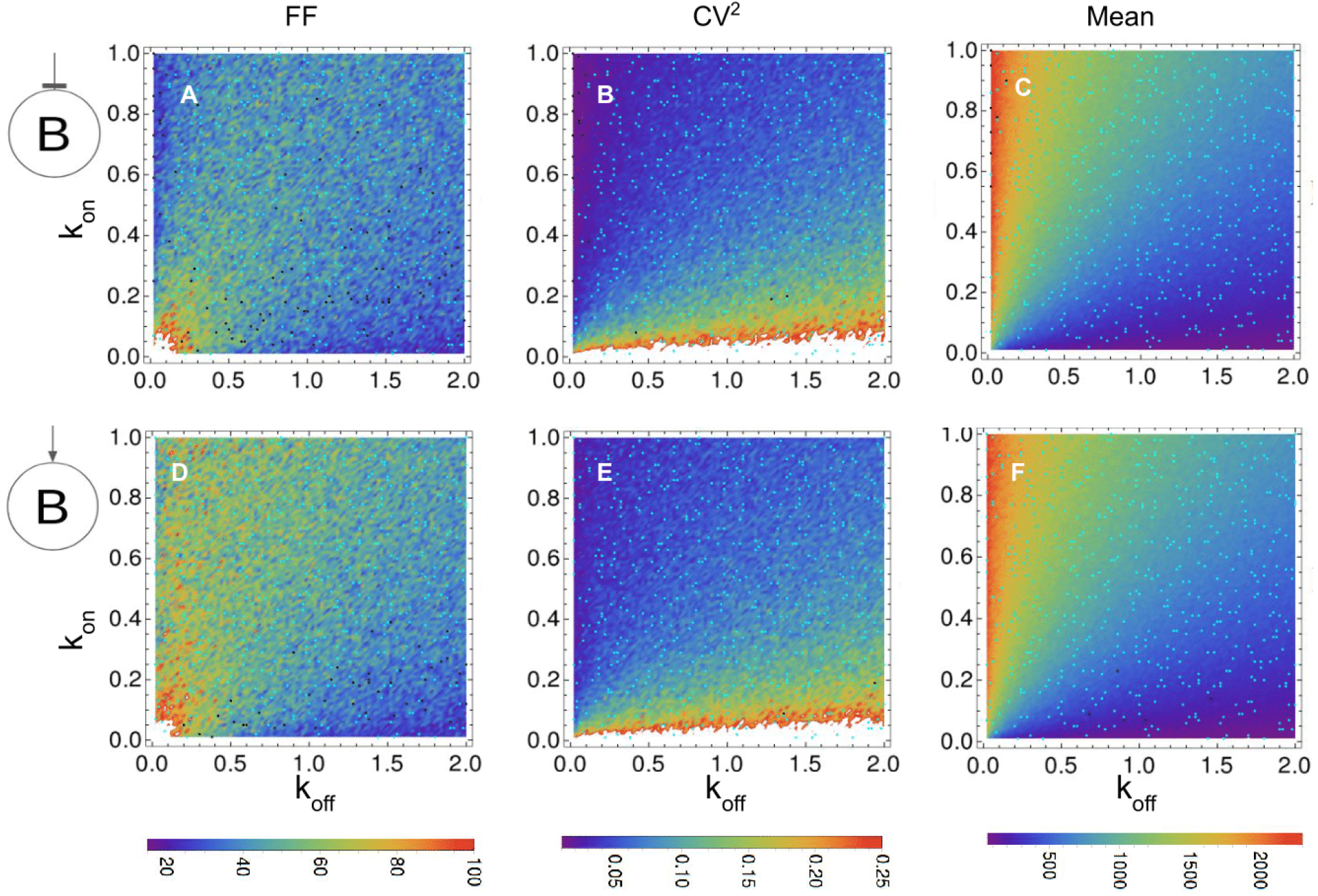
Estimates of FF, CV^2^, and mean of molecules at the protein level for inhibited and activated gene expression in a range of k_on_ and k_off_ parameter values. A-C. FF, CV^2^, and mean of molecules for an inhibited gene B, D-F. FF, CV^2^, and mean of molecules for an activated gene B. Here the parameters k_on_ and k_off_ of both regulated genes were changed while those for regulators remain unchanged. The black/cyan points indicate non-significant/significant differences, respectively, with an unregulated gene. The default parameters for both regulator and regulated genes are listed in Table S9. The regulator was expressed as in region 1. White spots are values higher than the scale plotted. Ccolor should be used for this figure in print.

An unregulated gene in region 3 had an almost continuous expression (Fig. 4A). However, it changed to a discontinuous expression when it was regulated by an activator with discontinuous burst expression (Fig. 4B). As a consequence, the variation in its expression was larger than that for an unregulated gene, which caused significant differences in FF values (Fig. 3D). On the contrary, in region 1, although the regulator also generated changes in the regulated gene (Fig. 4D), these changes were similar to those expected for an unregulated gene in this region (Fig. 4C). At the same time, when the regulated gene is expressed in burst as in region 1, the noise is propagated from regulator to regulated gene in an almost homogeneous manner (Fig. S4).

**Figure 4.**
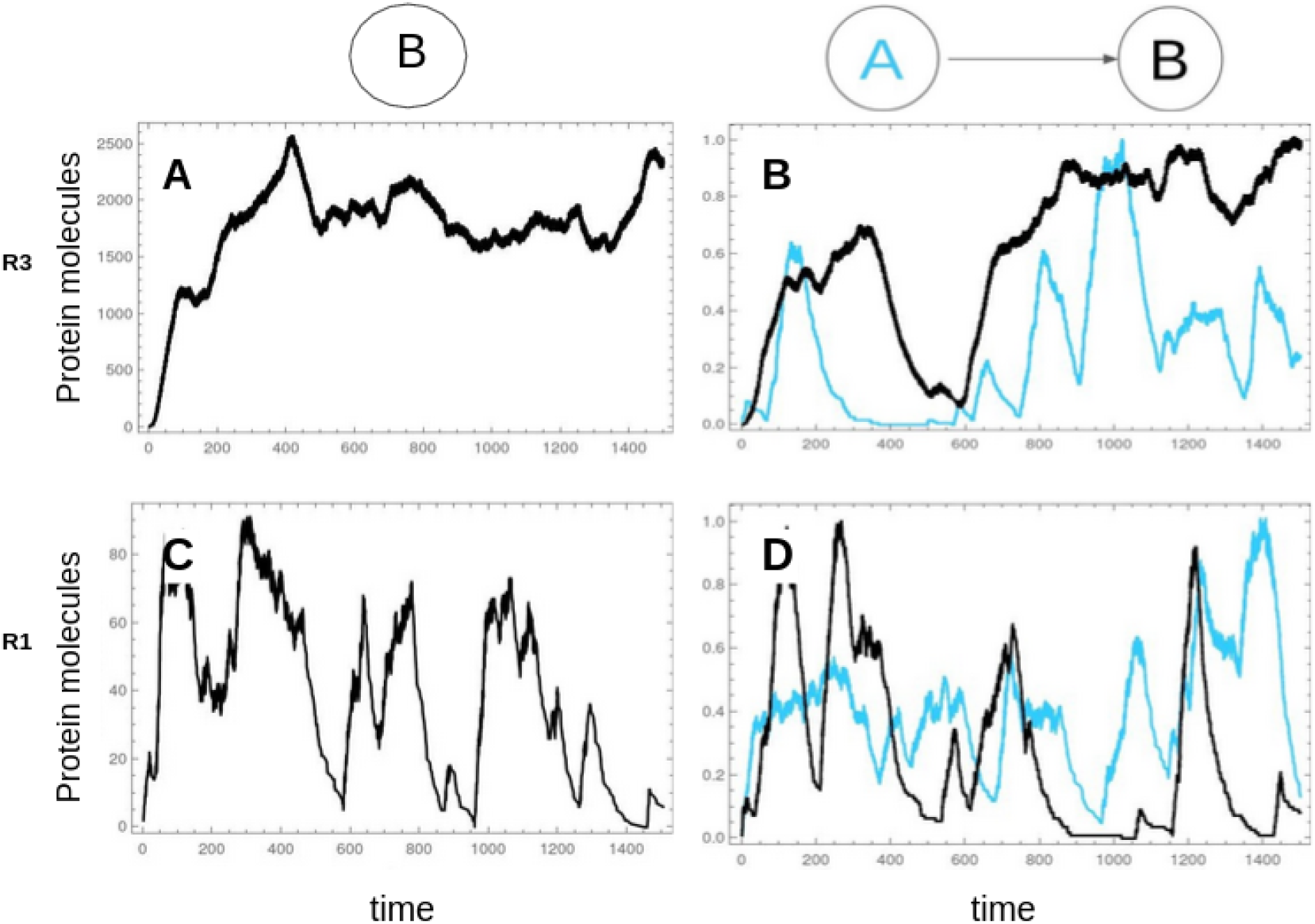
Temporal expression of a gene B in different regions with and without a regulator. A. Temporal expression of an unregulated gene B in region 3, B. Temporal expression of an activator A and an activated gene B, when B is expressed as in region 3, C. Temporal expression of an unregulated gene B in region 1, D. Temporal expression of an activator A and an activated gene B, when B is expressed as in region 1. The default parameters are listed in Table S9. Color should be used for this figure in print.

Figure S5 shows the case in which the promoter parameters were changed for the regulator but not for the regulated gene. Here, the inhibitor had an FF pattern similar to that for an unregulated gene (Fig.1A), but the inhibited gene was always expressed as in region 4. In region 4, when both the regulator and the regulated gene were expressed in a continuous way, the inhibition occurred with low values of both FF and mean of molecules (Fig. S5 A-B). In regions 3 and 2, when the regulator was expressed in large bursts and the regulated gene was expressed in a continuous way, the inhibition was lower than in previous region. This happened because sometimes the inhibitor had a low expression, which increased the FF and mean of molecules of the inhibited gene (Fig. S5 A-B). Finally in region 1, when the regulator was expressed in small bursts and the regulated gene was expressed in a continuous way, the inhibition didn’t occur and there were low FFs values and a high mean of molecules (Fig. S5 A-B).

### 3.4 Noise and expression level of the regulatory systems for coupled cells

It was possible to identify the four regions of expression for a self-activated gene simulated with the CLE (Fig. 5A-B). The noise (FF) pattern in a unicellular approach and throughout the range of kinetic parameters (i.e. k_off_, k_off_, kd_mRNA_, and kd_p_) was similar to that of a multicellular approach (Fig. 5 A-B, D-E, G-H). But the FF estimates for a multicellular approach were lower than for a unicellular approach (Fig. 5 A-B, D-E, G-H). In this multicellular system, the cells were coupled by diffusion of the self-activated gene product. This represents a paracrine signal that diffuses in the tissue and activates its expression in the cells expressing it.

**Figure 5.**
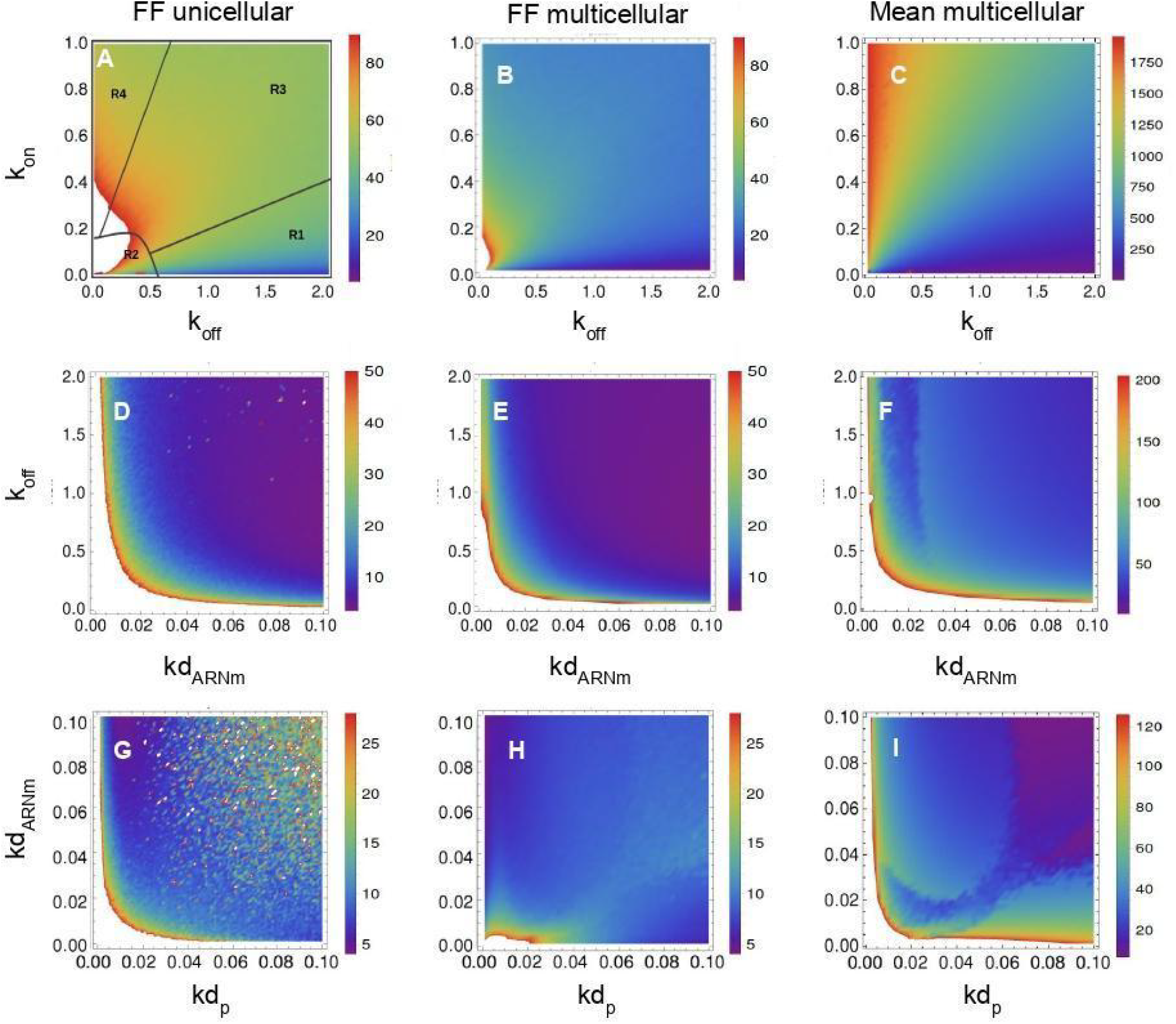
FF and mean of self-activated protein expression in a unicellular and multicellular approach for several kinetic parameters. A-C. Promoter activation k_on_ *vs* promoter inactivation k_off_, D-F. Promoter inactivation k_off_ *vs* mRNA degradation kd_mRNA_, G-I. mRNA degradation kd_mRNA_ vs protein degradacion kd_p_. Simulated with CLE. The default parameters are listed in Table S9. According to the parameters, the system is by default in region 1. Color should be used for this figure in print.

In a multicellular approach and throughout the region of k_off_ and mRNA degradation rate (kd_mRNA_), the noise (FF) and mean of molecules presented the highest values when both parameters were low, or when one of them was low and the other one fluctuated between low and high values (Fig.5 E-F). When both k_off_ and kd_mRNA_ were low, gene expression was as in region 2 (i.e., large bursts with high FF values). The increase in k_off_ returned the system to region 1 (i.e., small bursts with low FF values). However, the system remained in region 2 when kd_mRNA_ was very small (Fig.5 E-F).

In a multicellular approach throughout the range of kd_mRNA_ and kd_p_, the FF pattern had the highest values when both parameters were low (Fig.5 H). Therefore, the expression became more similar to that of region 2. At the same time, the mean of molecules presented the highest values when both were low (Fig. 5 I). The mean of molecules was also high when one of the parameters was low and the other one fluctuated between low and high values (Fig. 5 I).

For region 2-4, the FF decreased with the increase in diffusion rate (Fig. S6 A-D). Diffusion homogenized the expression levels of self-activated genes while preventing some cells from having high expression levels (Fig.6 D,F). There were some colony sizes for which the increase in diffusion rate increased the FF, which could indicate higher heterogeneity, but the colony actually had a more homogeneous expression than for lower diffusion rate values (Fig. 6 A,F). These high values of FF were due to differences in expression between the border cells and the rest of the colony (Fig.6 A,F). This resulted in a higher FF value for the whole colony than for a set of cells inside the colony (Fig. S7 A-B). At the same time, the increase in colony size decreased the FF (Fig.S6 A-D). But for regions 2-3, this happened after a threshold of minimum diffusion was reached (Fig.S6 B-C).

**Figure 6.**
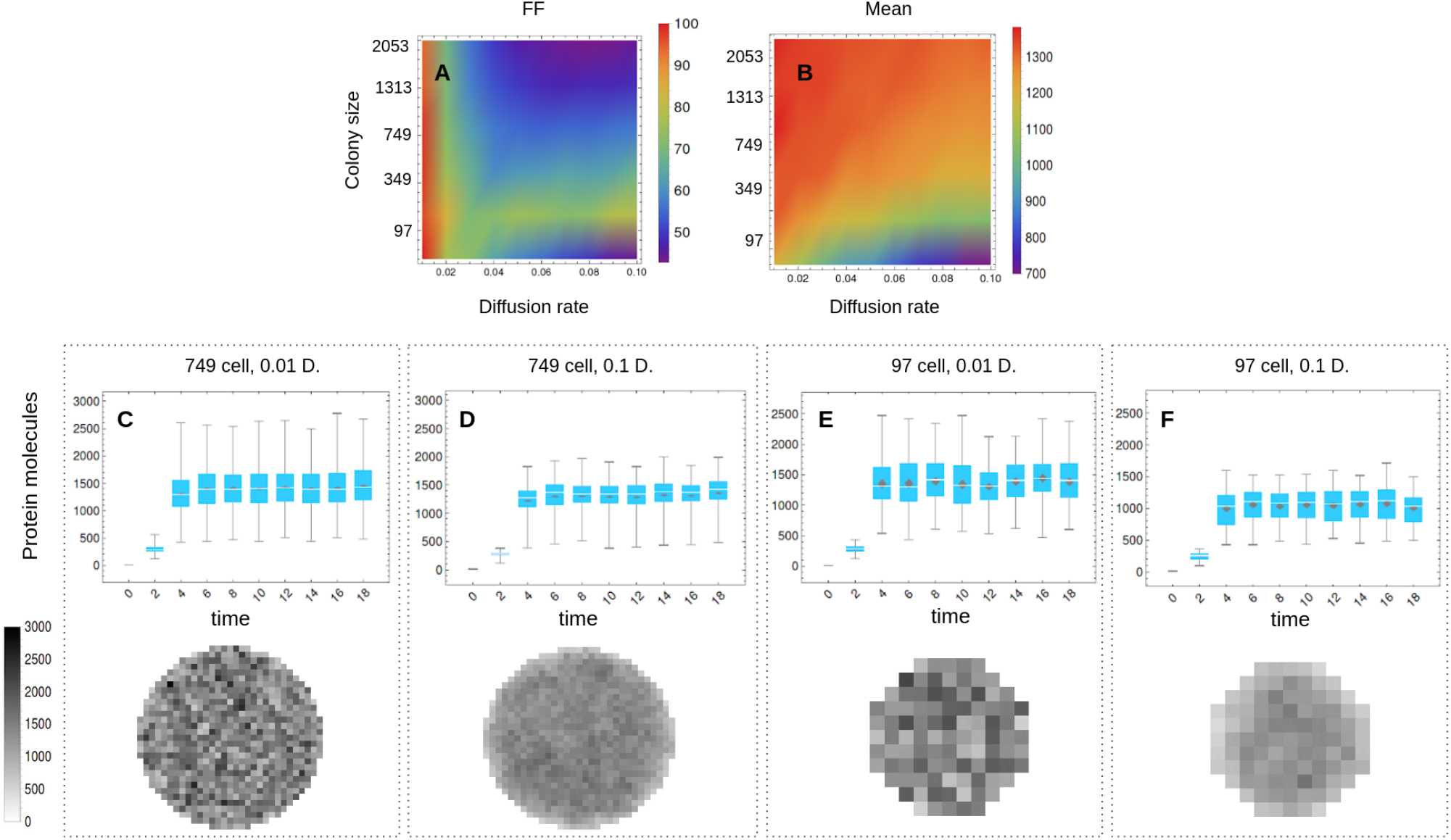
Estimates of FF and mean of molecules at the protein level for the expression of a self-activated gene in a cell colony and in a range of colony sizes and diffusion rate values for region 2. A-B. FF and mean of molecules of a self-activated gene with expression as in region 2, C-F. Top: Boxplots of protein molecules in the whole colony throughout time, Bottom: One state in time of the protein molecules in the cell colony. The default parameters for both regulator and regulated genes are in Table S9. Color should be used for this figure in print.

For regions 2-4, the mean of molecules decreased with the increase in diffusion rate but increased as colony size increased (Fig. S6 A-D). At the same time, the magnitude of the diminution in the mean as a consequence of diffusion depended on the colony size, with larger colonies having lower differences than smaller colonies (Fig.6 C-D and E-F). Finally, region 1 had a different behavior compared to the previously described regions, both in FF and mean of molecules. In this region, diffusion decreased FF estimates and there were no colony sizes for which it did not happen (Fig.S6 A and S7C-D), at the same time diffusion increased the mean of molecules (Fig. S6E).

## 4. DISCUSSION

### 4.1. Noise and expression levels in an unregulated gene

Although CV^2^ is considered the most direct and unambiguous measure of noise, FF reveals trends in noise that cannot be observed with CV^2^ when the molecule number is large [36]. The difference between them is that they measure different sources of noise; CV^2^ measures the noise due to the system size as it is explained by the *finite number phenomen* [6], and FF measures the noise due to promoter kinetics as it is observed in the SPK region. From previous studies, it is well known that SPK is the result of the chromatin remodeling process and it has an important role generating stochasticity in eukaryotic gene expression [17,47]. In fact, although it has been reported that genes with characteristics associated to transcriptional bursting are those with low values of *k*_*on*_ and relatively high values of *k*_*off*_ [17], due to SKP transcriptional bursting is present even when both *k*_*on*_ and *k*_*off*_ are equal or when *k*_*on*_ is larger than *k*_*off*_ whenever both have low values in the SKP region [10]. Because of these differences between both noise estimators, some studies indicate that CV^2^ measures the noise and FF measures the noise strength [36].

This study extends the previous analysis about the relationship between mean number of molecules, noise (CV^2^), and noise strength (FF). For instance, it is known that increasing gene expression through high *k*_*on*_ rates decreases noise and noise strength [6]. This happens because high *k*_*on*_ rates increase the mean of molecules and direct a more continuous expression [14]. On the other hand, increasing gene expression through lengthening of burst duration (i.e., decreasing *k*_*off*_ rate) is associated with more noise strength [22]. But, according to this study, this happens only when *k*_*on*_ values are low. Decreasing gene expression due to increases in *k*_*off*_ increases CV^2^ when the system has the largest mean values. This is followed by a constant CV^2^ value that is independent of the *k*_*off*_ value [36]. But again, according to this study, this happens only for some *k*_*on*_ values.

In this study, we identify four types of gene expression according to their dynamical behavior throughout the range of k_on_ and k_off_ values at the mRNA and protein levels. The dynamical behavior throughout these regions has different noise levels, noise strength, and expression levels. Throughout the regions gene expression went from happening in discontinuous bursts to being more continuous both at the mRNA and protein levels. With this classification it is possible to support facts such as that strong enhancers generate more bursts than weaker enhancers [37]. For instance, with strong enhancers k_on_ is larger than k_off_, which generates discontinuous bursts with high FF values, but only in region 2 (i.e., region with SPK), where there is an area with high burst sizes and mean of molecules.

It is well known that developmental gene expression is noisy, and this is due to transcriptional bursting [37] and low copy numbers of molecules [48]. Developmental genes such as *ush* and *hnt* are expressed in levels of around 200 molecules [14]. Other genes such as the Drosophila gap gene Kruppel has a CV^2^ of around 0.25 [21]. Therefore, because of number of molecules, the value of CV^2^, and the promoter activation and deactivation rates estimated for some developmental genes, we propose that those genes could have an expression similar to that seen in regions 1 and 2.

The FF value cannot be a measure to indicate whether or not there is regulation at the promoter level as previously pointed out in [8]. This is because at the protein level the FF value is higher than 1 in region 4, where expression is similar to a constitutive expression. Additionally, the FF can not be a measure to indicate whether or not there is discontinuous burst expression. This is because in both regions with expression in bursts (i.e., regions 1-2) the FF value is very different. Indeed, an expression in bursts could have a value of FF equal or lower than the FF for a more continuous expression. This means that the burst expression of genes does not necessarily imply high noise strength (FF).

### 4.2. Noise and expression levels of the regulatory systems for individual cells

Transcriptional bursts are an important feature of gene activity in living embryos [2]. But how do organisms establish and maintain the precise levels of gene expression seen during development? Some of the mechanisms described in literature include redundancy in genetic circuits to achieve the precision required for proper development [21]. It has also been reported that shadow enhancers filter the noise in TF due to input separation (i.e., each enhancer responds to a different TF). In this way, the target gene is less sensitive to levels of a single TF [21].

Other studies concluded that some GRNs, such as interlinked Feed-forward loops, are effective in filtering noise [16]. But there are different studies pointing out to different conclusions about self-activated genes. For instance in [16], feedback loops are noise controllers. On the contrary [22] indicates that when HIV becomes integrated into host genomes, it encodes its own positive transcriptional transactivator, Tat, and there is evidence of Tat-mediated amplification of basal HIV transcriptional noise. Other studies also argue that self-activation systems amplify fluctuations and the heterogeneity of populations [15,49,50]. In our study the self-activated gene did not have higher or lower noise levels than those of an unregulated gene. These discrepancies could be due to the model assumptions. For example, in [22], Tat protein regulation increases the noise strength when the protein affects either the transcriptional rate or the promoter deactivation rate; however, it decreases noise strength when it affects the promoter activation rate. In our model, the feedback protein affects the promoter activation rate but it does not generate significant changes in noise strength with respect to an unregulated gene.

Activator-inhibitor regulatory systems do not reduce the noise in gene expression when compared to an unregulated gene. On the contrary, noise levels are higher than those for an unregulated gene. These fluctuations in the expression of both genes involved in the activator-inhibitor regulatory system (i.e., activator and inhibitor) could thus facilitate its function in development. For instance, these fluctuations allow a heterogeneous expression of genes in time and space, which is necessary for self-organized fate patterning [45]. This means that noise could be beneficial in driving phenotypic diversity as previously pointed out in [22]. This increase in noise in both genes is due to noise propagation from the regulator to the regulated gene, as it will be explained in the next section.

### 4.3. Noise propagation from the regulator to the regulated gene

Previous studies concluded that fluctuations in regulatory proteins (i.e., TFs and signaling molecules) propagate down a GRN [21]. This significantly alters the expression and noise levels of downstream target genes [21]. Specifically, the propagation of noise is associated with a reduction in the mean of molecules. In this study, we propose new and important ideas about noise propagation.

First, the noise propagation depends on the expression type (i.e. four regions of expression) of both the regulator and the regulated gene. For instance, if the regulator is expressed in bursts (i.e. regions 1-2) there is a significant increase in noise if the regulated gene is expressed as in region 3-4. If, on the other hand, the regulated gene is expressed in bursts, the noise propagates through a regulatory connection independently of the amount of noise of the regulator, and sometimes there is no significant increase in noise.

This dependency on expression type explains why the noise in the self-activated system is not higher than in an unregulated system. This is because, for this case, both the regulator and the regulated gene, which are the same gene, have the same expression type. Therefore, according to this study, self-activation does not increase noise but activation by another gene does.

The second important aspect of noise propagation is that it depends on the type of regulation (i.e. activation and inactivation). As previously mentioned in [1,51], the stochasticity and low intracellular copy number limit the cellular signal precision and then the gene regulation. This is the reason behind the differences in noise propagation between activation and inactivation. For instance, inhibition is less effective when the inhibitor is expressed in bursts (i.e. region 1) and the regulated gene’s expression is as in region 4. Thus, the regulated gene’s expression is independent of regulator expression. But this does not happen for activation.

### 4.4. Noise and expression levels of regulatory systems for coupled cells

The noise strength for the self-activated system in a cell colony also varies throughout the promoter activation and inactivation rate range. Additionally, it also depends on transcriptional and translational efficiency (i.e., ks_mRNA_/k_off_ and ks_p_/kd_mRNA_ respectively). For instance, the increase in transcriptional and translational efficiency increases fluctuations in protein dynamics [22][31]. This means that both at the unicellular and multicellular levels, the kinetic parameters are modulators of noise. Not only are the kinetic parameters associated with gene expression, but diffusion and colony size also affect the heterogeneity and magnitude of expression levels of a self-activated gene. In fact, both diffusion and colony size are noise controllers as it will be explained below. This is very important to understand the community effect in embryonic development.

The community effect happens when cells in a tissue are coordinated to express the same set of genes that specify a differentiation phenotype [23,24]. All cells in the community express the same set of target genes because a paracrine signal activates its own expression in receptor cells as well as the expression of target genes [52]. This role is clearly represented by the self-activated gene simulated here. A minimum number of cells is required for the effect to occur; below this level,cells could not differentiate [24]. For instance, in the induction of tooth formation a minimum of 25% of inductive tooth mesenchyme cells in the tissue is necessary [52]. In other words, the diffusion of a paracrine signal throughout a colony of cells decreases noise.

Some previous studies have modeled this phenomenon and they have included the promoter regulation in their models [23,24]. But none of them have evaluated the community effect with different types of expression. Additionally, none of them have evaluated the effect of the rate of diffusion as evaluated here. For instance, in [24] a cellular communication parameter is evaluated, but it is not analogous to our diffusion rate because the increase in their parameter increases the mean of molecules, while our parameter decreases it. Here we propose that minimum colony sizes are required to achieve the community effect because larger cell colonies counteract diffusion losses and diffusion is necessary to decrease noise and synchronize gene expression across cells. But this only applies to gene expression in regions 2-4. This homogenization of signal levels throughout the colony is important to send a more robust signal to target genes.

## 5. CONCLUSION

The noise levels in self-activated and activator-inhibitor regulatory systems depend on the gene expression type of both the regulator and the regulated gene. In this way, the particular forms in which genes connect to each other in these regulatory systems do not explain the noise in expression. However, the noise has a propagation pattern different for activation and inactivation types of regulation. The changes in kinetic parameters affect the noise levels; in this way, all mechanisms that change their values could be potential noise filtering mechanisms. Similarly, diffusion and colony size could be mechanisms of noise filtering in gene expression in a colony of cells. The increase in diffusion rate and colony size are necessary to synchronize gene expression and perform the community effect.

## 6. ACKNOWLEDGEMENTS

We like to thank Jayson Gutierrez and Tatiana Rodriguez for their advices. We thank Amalia Llano for his comments and criticisms of the first draft. All authors acknowledge partial support from CODI-Universidad de Antioquia under project “Estudio de la Evolución de Circuitos de Regulación del Desarollo Embrionario a través de la Biología Evolutiva de Sistemas” code 2017-14367. The funders had no role in study design, data collection and analysis, decision to publish, or preparation of the manuscript.

