## Supplemental Tables and Figures for "Study and Modeling of Biological Noise-Filtering Properties of Conserved Gene Regulatory Networks Motifs in Animal Development"

| Parameter | Symbol | Unity |
|---|---|---|
| Promoter activation rate | $k_{on}$ | *1/min* |
| Promoter inactivation rate | $k_{off}$ | *1/min* |
| Transcription rate | $ks_{mRNA}$ | *mRNAs/min* |
| mRNA degradation rate | $kd_{mRNA}$ | *mRNAs/min* |
| Translation rate | $ks_{p}$ | *protein/min* |
| Protein degradation rate | $kd_{p}$ | *protein/min* |
| Diffusion rate | $D$ | *pix$^{2°}$/min* |

**Table S1.** ***Symbols of the parameters used in this study.***

| kon value | Koff value | Gen/cell type | The function of the gene | Ref. |
|---|---|---|---|---|
| 0.023-0.033 | 0.26-0.35 | *Bmal1a* promoter in Mouse fibroblast single-cell | *It generates molecular circadian rhythms and it* is the only clock gene without which the circadian clock fails to function in humans *(wiki)* | [1] |
| 0.018-0.032 | 0.16-0.28 | *Glutaminase* promoter in Mouse fibroblast single-cell | *It generates glutamate from glutamine.Glutamate is the most abundantly used excitatory neurotransmitter in the CNS (Wikipedia)* | [1] |
| 0.0225-0.06 | 0.48-0.58 | *Plectin1* promoter of Mouse fibroblast single-cell | *It is implied in maintaining cell and tissue integrity, and it is a scaffolding platform for the assembly, positioning, and regulation of signaling complexes (https://www.ncbi.nlm.nih.gov/gene/5339)* | [1] |
| 0.023-0.045 | 0.22-0.36 | *Serpine1* promoter of Mouse fibroblast single-cell | *It is an inhibitor of fibrinolysis and also functions as a component of innate antiviral immunity (https://www.ncbi.nlm.nih.gov/gene/5054)* | [1] |
| 0.021-0.047 | 0.19-0.27 | *Sh3kbp1* promoter of Mouse fibroblast single-cell | *It facilitates protein-protein interactions and has been implicated in numerous cellular processes including apoptosis, cytoskeletal rearrangement, cell adhesion, and in the regulation of clathrin-dependent endocytosis (https://www.ncbi.nlm.nih.gov/gene/30011)* | [1] |
| 0.018-0.032 | 0.1-0.15 | *Prl2C2* promoter of Mouse fibroblast single-cell | | [1] |
| 0.25 - 0.32 | 0.04 - 0.06 | *Ush gene promoter in Drosophila embryos at the onset of nuclear cleavage cycle 14 (nc14) with high BMP level* | *A gradient of bone morphogenetic protein (BMP) signaling patterns ectodermal cell fates along the dorsal-ventral axis of vertebrate and invertebrate embryos. The BMP/pMad gradient activates* | [2] |

| | | | | |
|---|---|---|---|---|
| | | | *different thresholds of gene activity, including the intermediate target u-shaped (ush)* | |
| 0.19- 0.28 | 0.04 - 0.06 | *Ush gene promoter in Drosophila embryos at the onset of nuclear cleavage cycle 14 (nc14) with medium BMP level* | --- | [2] |
| 0.07 - 1.7 | 0.04 - 0.06 | *Ush gene promoter in Drosophila embryos at the onset of nuclear cleavage cycle 14 (nc14) with low BMP level* | --- | [2] |
| 0.46 - 0.5 | 1.9- 2.1 | *hnt gene promoter in Drosophila embryos at the onset of nuclear cleavage cycle 14 (nc14) with high BMP level* | *A gradient of bone morphogenetic protein (BMP) signaling patterns ectodermal cell fates along the dorsal-ventral axis of vertebrate and invertebrate embryos. The BMP/pMad gradient activates different thresholds of gene activity, including the peak target gene hindsight (hnt)* | [2] |
| 0.3 - 0.4 | 1.9- 2.1 | *hnt gene promoter in Drosophila embryos at the onset of nuclear cleavage cycle 14 (nc14) with medium BMP level* | --- | [2] |
| 0,01158 | *2.082* | For none in specific | *Their simulation parameters are all within biologically or biochemically relevant range, which usually spans orders of magnitude, such as the one for protein degradation rates. Where possible, they have referred to reaction rates that have been determined experimentally to choose parameter values for simulations* | [3] |
| 0.045 | 0.005 | *Human cyclin D1 (CCND1) in HEK-293 cells* | *Different cyclins exhibit distinct expression and degradation patterns which contribute to the temporal coordination of each mitotic event.* | [4] |
| | *0.2 (0, 0.05)* | For none in specific | --- | [5][1] |
| 1.0 | *1.0* | NODAL | They indicate neither the dimensions nor the source of this date. It is an entirely theoric study. | [6] |
| 1.0 | *1.0* | LEFTY | They indicate neither the dimensions nor the source of this date. It is an entirely theoric study. They made a bifurcation diagram for $k_{on}$ parameter between 0-1.4 | [6] |

**Table S2.** ***Activation and Inactivation promoter rate*** *($k_{on}$ - 1/min, $k_{off}$ - 1/min)*

| Value | Gen/cell type | The function of the gene | Ref. |
|---|---|---|---|
| 0.1-0.28 | *Plectin1* promoter of Mouse fibroblast single-cell | --- | [1] |
| 0.2 - 0.3 | *Bmal1a* promoter in Mouse fibroblast single-cell | --- | [1] |
| 0.8 - 1.2 | *Serpine1* promoter of Mouse fibroblast single-cell | --- | [1] |
| 1.8 - 2.0 | *Sh3kbp1* promoter of Mouse fibroblast single-cell | --- | [1] |

| | | | |
|---|---|---|---|
| 2.0 - 3.0 | *Glutaminase* promoter in Mouse fibroblast single-cell | --- | [1] |
| 3.4 - 4.3 | *Prl2C2* promoter of Mouse fibroblast single-cell | --- | [1] |
| 0.31-0.78 kb/min | *Human cyclin D1 (CCND1) in HEK-293 cells* | --- | [4] |
| 9 - 15 Poll II / min (loading rate) | *Ush gene in Drosophila embryos at the onset of nuclear cleavage cycle 14 (nc14), it does not depend of BMP level* | --- | [2] |
| 37 - 45 Poll II / min (loading rate) | *hnt gene in Drosophila embryos at the onset of nuclear cleavage cycle 14 (nc14), it does not depend of BMP level* | --- | [2] |
| 0.696 | *Non-specific* | It is for a gene making part of the community effect subcircuit | *[3]* |

**Table S3.** ***Transcription rate*** *($ks_{mRNA}$ - mRNAs/min)*

| Value | Gen/cell type | The function of the gene | Ref. |
|---|---|---|---|
| 0,02082 | Non-specific | It is for a gene making part of the community effect subcircuit | [3] |
| 0.0083 | NODAL | --- | [7] |
| 0.0041 | LEFTY | --- | [7] |

**Table S4. mRNA degradation rate** ($kd_{mRNA}$ - mRNAs/min)

| Value | Gen/cell type | The function of the gene | Ref. |
|---|---|---|---|
| *23 - 29* | *Ush gene promoter in Drosophila embryos at the onset of nuclear cleavage cycle 14 (nc14) with high BMP level* | --- | [2] |
| *15 - 25* | *Ush gene promoter in Drosophila embryos at the onset of nuclear cleavage cycle 14 (nc14) with medium BMP level* | --- | [2] |
| *5 - 16* | *Ush gene promoter in Drosophila embryos at the onset of nuclear cleavage cycle 14 (nc14) with low BMP level* | --- | [2] |

**Table S5.** ***mRNA transcriptional burst size*** *($b_{mRNA} = ks_{mRNA} * T'on' = ks_{mRNA} / Koff$)*

| Value | Gen/cell type | Function of the gene | Ref. |
|---|---|---|---|
| 1.386 | Non specific | It is for a gene making part of the community effect subcircuit | [3] |
| 2 (proteins/min/mRNA) | Nodal | --- | [7] |

| | | | |
|---|---|---|---|
| 2 (proteins/min/mRNA) | Lefty | --- | [7] |

**Table S6.** ***Translation rate*** *($ks_p$ - protein/min)*

| Value | Gen/cell type | The function of the gene | Ref. |
|---|---|---|---|
| 0.02082 | Non-specific | It is for a gene making part of the community effect subcircuit | [3] |
| 0.1 | Nodal | They indicate neither the dimensions nor the source of this date. It is an entirely theoric study. | [6] |
| 0.1 | Lefty | They indicate neither the dimensions nor the source of this date. It is an entirely theoric study. | [6] |

**Table S7.** ***Protein degradation rate*** *($kd_p$ - protein/min)*

| Value | Gen/cell type | The function of the gene | Ref. |
|---|---|---|---|
| $0.7\pm0.2\ \mu m^2/s$ (Cyclops-GFP)<br>$3.2\pm0.5\ \mu m^2/s$ (Squint-GFP) | Nodal | --- | [8] |
| $11.1\pm0.6\ \mu m^2/s$ (Lefty1-GFP)<br>$18.9\pm3.0\ \mu m^2/s$ (Lefty2-GFP) | Lefty | --- | [8] |

**T*able S8. Extracellular Effective diffusion coefficients***

| Parameter | Symbol | Unity | Constitutive | Unregulated | Auto-activated | Activator-Inhibitor |
|---|---|---|---|---|---|---|
| Promoter activation rate | $k_{on}$ | 1/min | --- | 0,01158 | 0,01158 | 0,01158 |
| Promoter inactivation rate | $k_{off}$ | 1/min | --- | 2,08 | 2,08 | 2,08 |
| Transcription rate | $ks_{mRNA}$ | mRNA/ min | 0,696 | 0,696 | 0,696 | 0,696 |
| mRNA burst size | $b_m$ | mRNAs | --- | 0,333 | 0,333 | 0,333 |
| mRNA degradation rate | $kd_{mRNA}$ | mRNA/ min | 0,02082 | 0,02082 | 0,02082 | 0,02082 |
| Translation rate | $ks_p$ | protein/ min | 1,386 | 1,386 | 1,386 | 1,386 |
| Protein burst size | $b_p$ | protein/ mRNA | --- | 66,57 | 66,57 | 66,57 |
| Protein degradation rate | $kd_p$ | protein/ min | 0,02082 | 0,02082 | 0,02082 | 0,02082 |
| Auto-activation constant | $k_{AA}$ | protein | --- | --- | 100 | 100 |
| Inhibition constant | $k_{AI}$ | protein | --- | --- | --- | 100 |
| Inhibition constant | $k_{AI}$ | protein | --- | --- | --- | 100 |
| Hill constant | h | --- | --- | --- | 3 | 3 |
| Diffusion rate | *Da* | $pix^{2°}/min$ | --- | --- | 0.01 | ---- |

**T*able S9. Default parameters values of the regulatory systems evaluated in this study.*** *The majority of these were from [3].* These parameters apply both for activator (A) and inhibitor (I). ° 1 pixel ~ 16 µm. mRNA and protein are molecules.

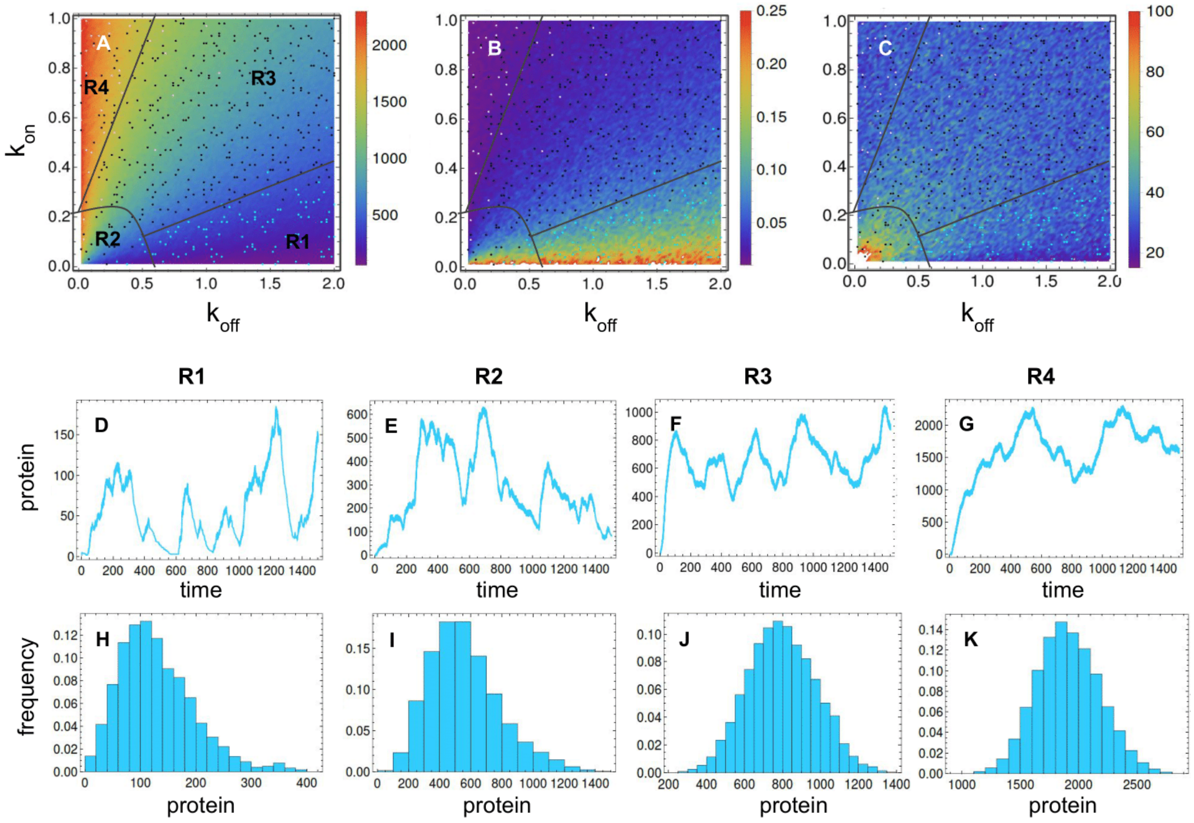


**Figure S1.** ***Regions of gene expression types at the protein level for an unregulated gene in a range of $k_{on}$ and $k_{off}$ parameter values.*** A. The four regions over the mean of molecules pattern B. The four regions over the CV² pattern, C. The four regions over the FF pattern, D-G. The temporal dynamics of expression in each region, H-K. The steady-sate distribution in each region. Default parameters are listed in Table S9. The color of points represents the clustering group to which the pair of parameters belongs to. Color should be used for this figure in print.

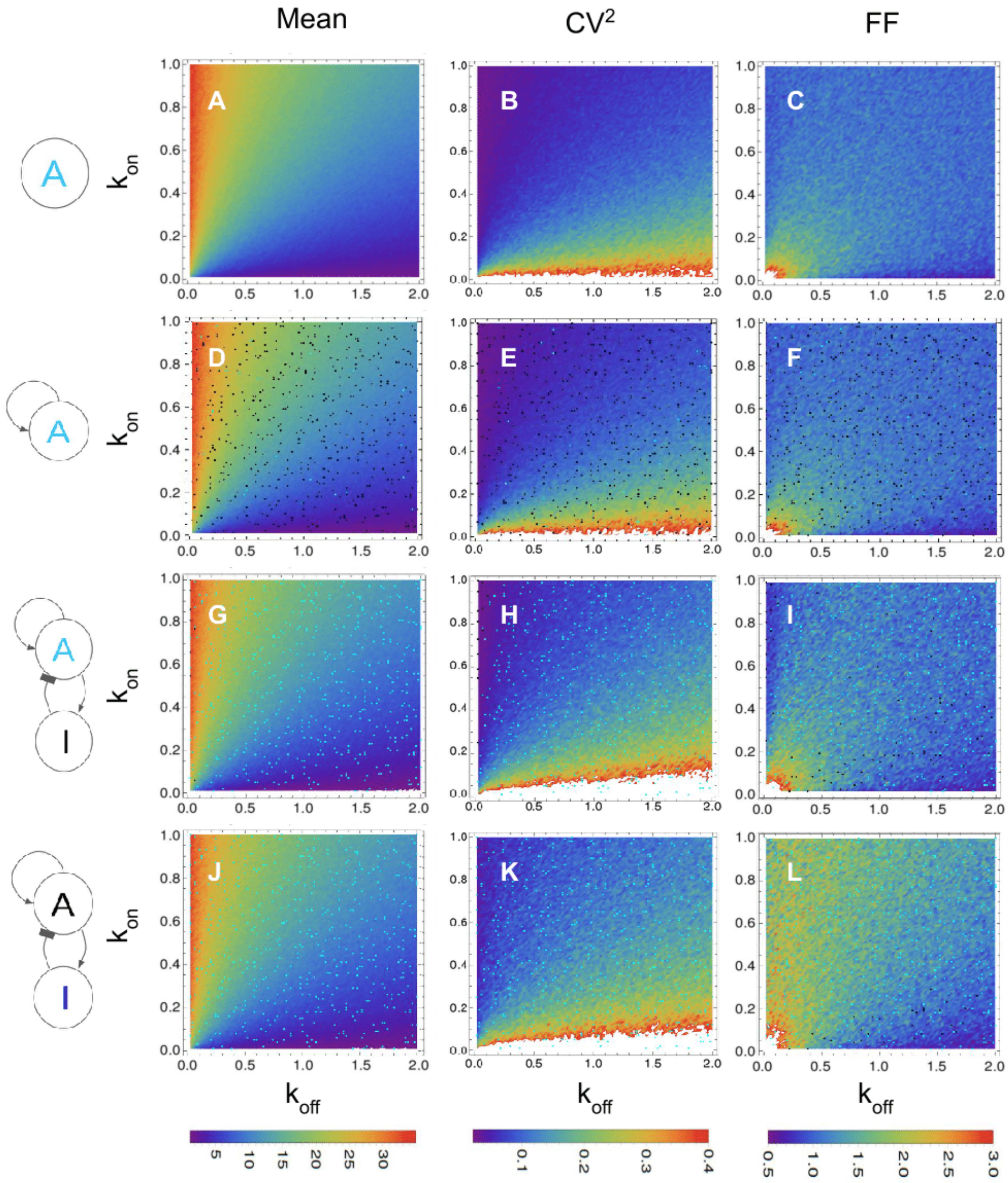


**Figure S2.** ***Estimates of mean of molecules, $CV^2$, and FF at the mRNA level for gene expression of the regulatory systems evaluated in a range of values for $k_{on}$ and $k_{off}$.*** A-C. mean, $CV^2$, and FF for an unregulated gene A, D-F. mean, $CV^2$, and FF for a self-activated gene A, G-I. mean, $CV^2$, and FF for gene A in an activator-inhibitor regulatory system when only the parameters of gene A were changed, J-L. mean, $CV^2$, and FF for gene I in an activator-inhibitor regulatory system when only the parameters of gene I were changed. The black/cyan points indicate non-significant/significant differences when compared to an unregulated gene. The default parameters are found in Table S9. White points are values higher than the scale plotted. Color should be used for this figure in print.

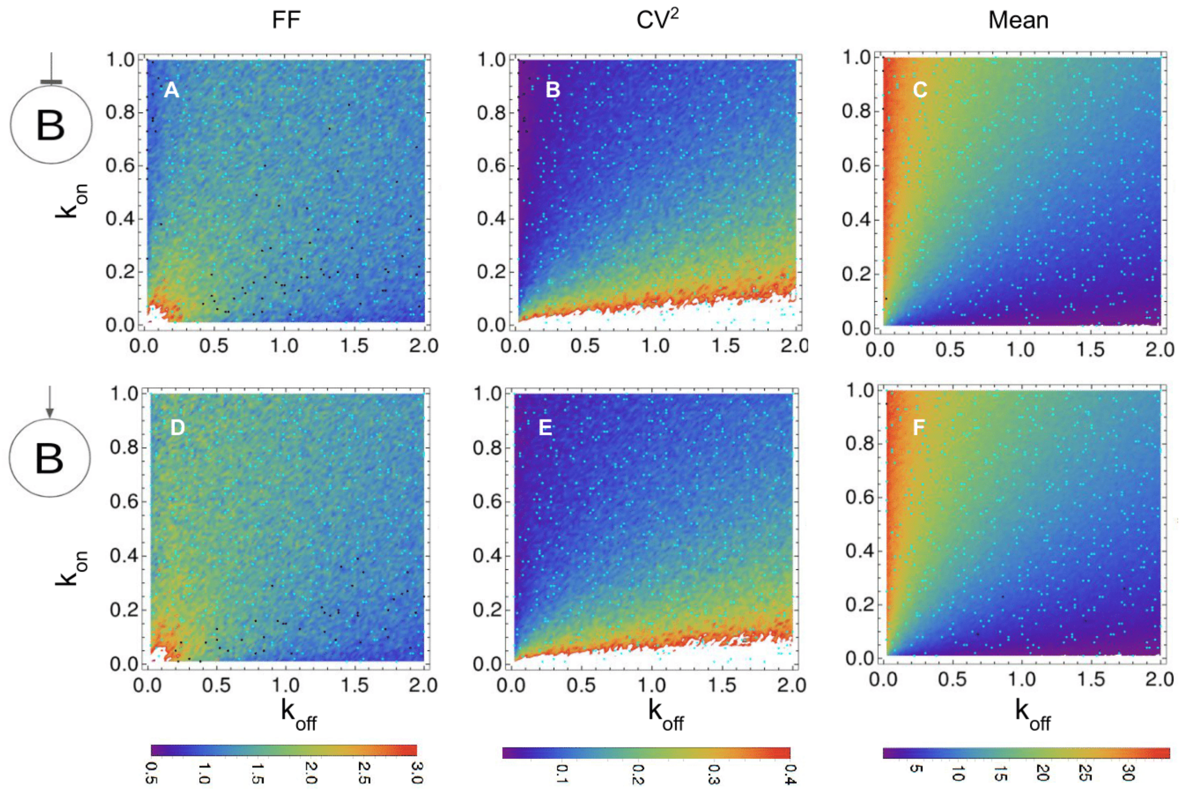


**Figure S3.** ***Estimates of FF, CV$^2$, and mean of molecules at the mRNA level for inhibited and activated gene expression in a range of $k_{on}$ and $k_{off}$ parameter values.*** A-C. FF, CV$^2$, and mean of molecules for an inhibited gene B, D-F. FF, CV$^2$, and mean of molecules for an activated gene B. Here the parameters $k_{on}$ and $k_{off}$ of both regulated genes were changed while those for regulators remain unchanged. The black/cyan points indicate non-significant/significant differences, respectively, with an unregulated gene. The default parameters for both regulator and regulated genes are listed in Table S9. The regulator was expressed as in region 1. White points are values higher than the scale plotted. Color should be used for this figure in print.

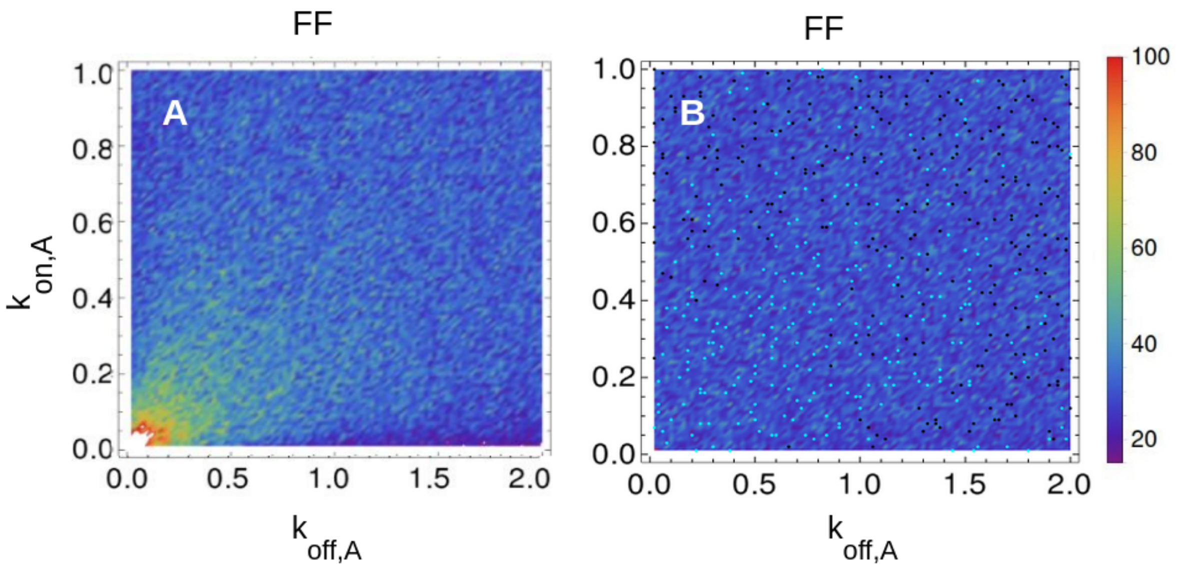


**Figure S4.** ***Estimates of FF at the protein level for regulator and regulated gene in a range of $k_{on}$ and $k_{off}$ parameter values.*** A. FF for activator when the values of its $k_{on}$ and $k_{off}$ parameters are changed, B. FF for activated gene whenwhen the values of its $k_{on}$ and $k_{off}$ parameters are fixed. The black/cyan points indicate non-significative/significative differences an unregulated gene. The regulated gene is expressed as in region 1. The default parameters for both regulator and regulated genes are listed in Table S9. White points are values higher than the scale plotted. Color should be used for this figure in print.

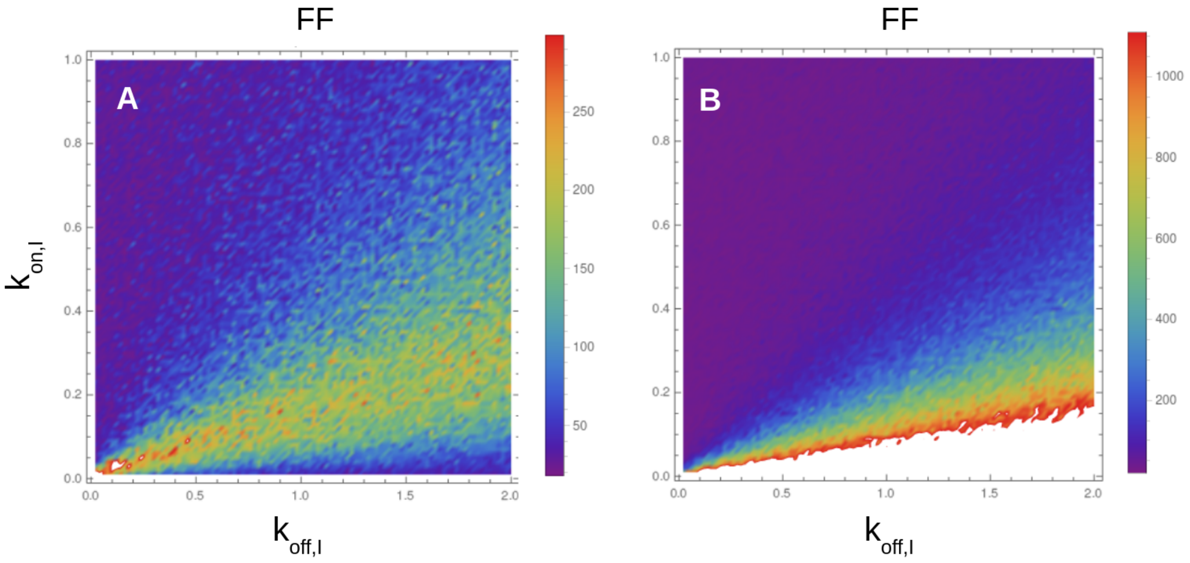


**Figure S5.** ***Estimates of FF and mean of molecules at the protein level for an inhibited gene in a range of $k_{on}$ and $k_{off}$ parameter values.*** A. FF for an inhibited gen when the values of its $k_{on}$ and $k_{off}$ parameters are fixed but those of its inhibitor are changed, B. Mean for inhibited gen when the values of its $k_{on}$ and $k_{off}$ parameters are fixed but those of its inhibitor are changed. The black/cyan points indicate non-significative/significative differences with an unregulated gene. The regulated gene is expressed as in region 4. The default parameters for both regulator and regulated genes are listed in Table S9. White points are values higher than the scale plotted. Color should be used for this figure in print.

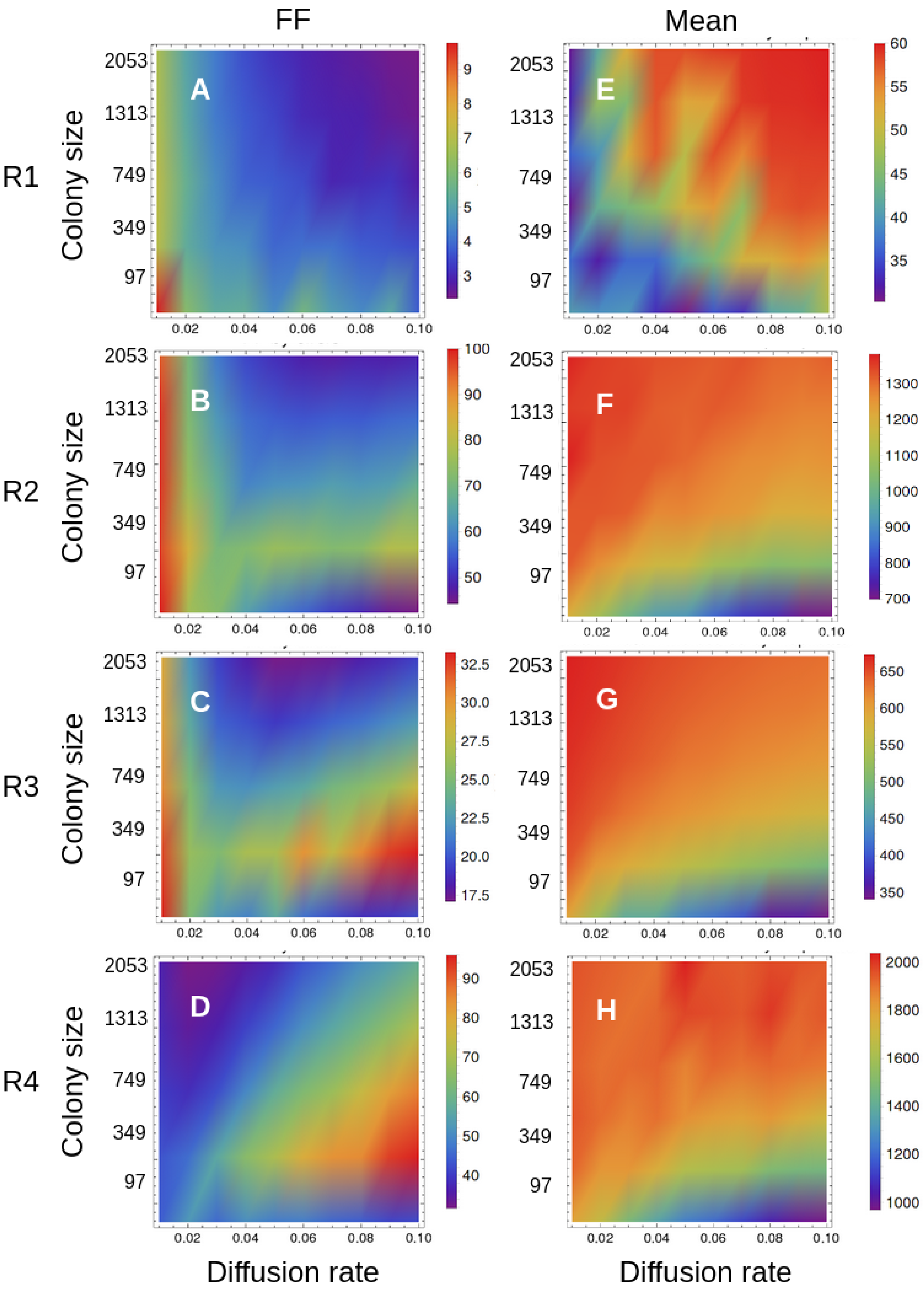


**Figure S6.** ***Estimates of FF and mean of molecules at the protein level for a self-activated gene in a cell colony for different values of colony sizes and diffusion rates.*** A-B. FF and mean of molecules when self-activated gene expresses as in region 1, C-D. FF and mean of molecules when self-activated gene expresses as in region 2, E-F. FF and mean of molecules when self-activated gene expresses as in region 3, G-H. FF and mean of molecules when self-activated gene expresses as in region 4. The default parameters are listed in Table S9. Color should be used for this figure in print.

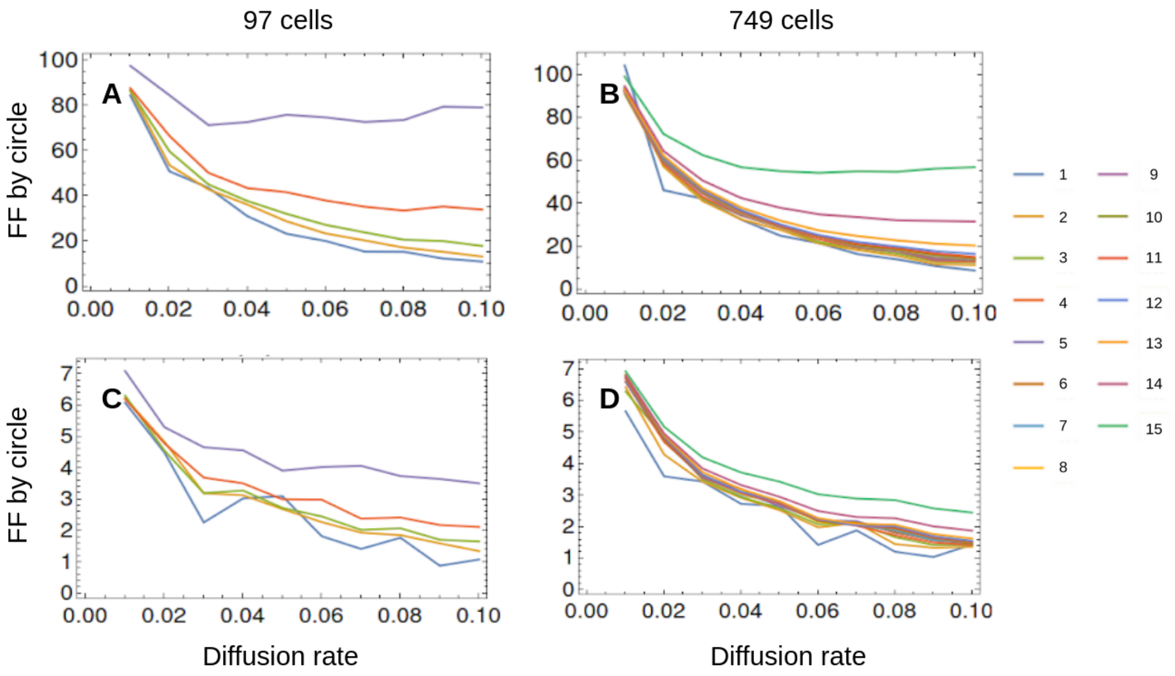


**Figure S7.** ***Estimates of FF at the protein level for a self-activated gene in a colony.*** *In each plot, each line represents the changes in FF throughput the diffusion rate for a set of cells in the colony. Each set of cells are forming concentric circles from the center to the outside of the colony.* A. For a colony of 21 cells and a self-activated gene with expression as in region 1, B. For a colony of 749 cells and a self-activated gene with expression as in region 1, C. For a colony of 21 cells and a self-activated gene with expression as in region 2, D. For a colony of 749 cells and a self-activated gene with expression as in region 2. The default parameters are listed in Table S9. White points are values higher than the scale plotted. Color should be used for this figure in print.
